# Understanding the dynamics of physical activity practice in the health context through Regulatory Focus and Self-Determination theories

**DOI:** 10.1101/623504

**Authors:** Manon Laroche, Peggy Roussel, François Cury, Julie Boiche

## Abstract

This research aimed to associate for the first time in the literature the Regulatory Focus and Self-Determination theories to understand the dynamics of physical activity practice in the health context. Two cross-sectional studies were conducted with 603 (Study 1) and 395 (Study 2) French volunteer participants aged from 18 to 69 and 19 to 71 respectively, and healthy or concerned by a health condition. The main results of structural equation modeling analyses demonstrated that across the two studies, health promotion focus was positively associated with intrinsic motivation (.44 < β < .74, p < .001), integrated regulation (.47 < β < .72, p < .001), identified regulation (.40 < β < .69, p < .001) and introjected regulation (.41 < β < .53, p < .001), whereas health prevention focus was positively related with external regulation (.31 < β < .45, p < .001) and amotivation (.32 < β < .38, p < .001). Bootstrapping analyses main results in Study 2 showed that health promotion focus was indirectly associated with physical activity through intrinsic motivation (95% CI [.02 to .11]), integrated regulation (95% CI [.00 to .08]), identified regulation (95% CI [.00 to .09]) and introjected regulation (95% CI [.04 to .12]), whereas health prevention focus was indirectly associated with physical activity through external regulation (95% CI [.00 to .12]). These studies reveal meaningful associations between Regulatory Focus and Self-Determination theories’ variables which support the relevance of associating these two models to understand the processes underlying the physical activity practice.

## INTRODUCTION

In recent years, the consequences of a lack of Physical Activity (PA), both for individuals’ health and in terms of costs for health systems [1] have led governments and health professionals to wonder about their capacity to modify people’s lifestyles through various PA promotion strategies. The Global Action Plan adopted by the World Health Organization aims for example to reduce the lack of PA by 10% by 2025 [2]. This new awareness is also manifested in the development of new technologies favoring PA and in particular by the development on the market of coaching apps which greatly facilitate access to the practice. However, while health professionals and public health communication campaigns recommend being physically active on a regular basis, almost half of all Europeans do not practice PA and this proportion has grown steadily in recent years [3]. A better understanding of motivational issues surrounding engagement in PA thus becomes of utmost importance.

Based on the Regulatory Focus Theory (RFT, [4]), previous studies showed that *promotion* and *prevention* regulatory foci are two significant motivational determinants of the adoption of various health behaviors [5]. Promotion focus involves a strategic inclination to be enthusiastic by approaching matches with the desired end-states, while prevention focus involves a strategic inclination to be vigilant by avoiding mismatches with the desired end-states [6]. In the health context, promotion focus refers to a chronic tendency to seek health-related gains and opportunities for improvement of one’s health state, while health prevention focus refers to a chronic tendency to avoid health-related losses and threats that could harm the maintenance of one’s health state [7, 8, 9].

To date, only one study [10] has examined the links between these chronic Health Regulatory Foci (HRF) and PA practice. In particular, these authors have evidenced among sport practitioners that health promotion focus was positively associated with amount of sports practice whereas health prevention focus was negatively related with this variable. But the issue of PA is not limited to a population of sport practitioners. Thus, the understanding of the links between HRF and PA should be extended to a wider population, taking into account a less specific PA than sport practice. Furthermore, Laroche et al. [10] evidenced one relevant process (i.e., Selection, Optimization and Compensation [SOC] strategy) which explained the positive link between health promotion focus and amount of sports practice. However, this psychological process was not found to be appropriate to explain the negative link between health prevention focus and amount of sports practice. To improve understanding of the relationship between HRF and PA, other psychological processes could undoubtedly be operative and would call for empirical attention.

Self-Determination Theory (SDT, [11]) has become a popular support for understanding motivation for PA, in particular in the health context [12]. This model distinguishes *intrinsic motivation* (PA is performed for the pleasure and satisfaction directly derived from it), *extrinsic motivation* (PA is performed to obtain outcomes that are distinct from the activity itself) and *amotivation* (individuals feel that there are no positive outcomes to be expected from PA). Extrinsic motivation refers to four types of regulation reflecting various levels of self-determination: *integrated regulation* (when PA is seen as congruent with individuals’ core values and lifestyle), *identified regulation* (when PA enables attaining a personally valued goal), *introjected regulation* (when individuals practice PA to avoid negative feelings of shame or guilt, or to enhance feelings of self-worth) and *external regulation* (when PA is controlled by external contingencies). There is an abundant literature linking motivation as conceived by SDT and PA, both in healthy adults and in individuals concerned by a health condition. Intrinsic motivation, integrated regulation and identified regulation were consistently found to be positively associated with PA indicators; null or positive associations were found for introjected regulation, while external regulation and amotivation mostly showed non-significant or negative relationships [13]. The more or less self-determined motivation developed toward PA could thus be a plausible process candidate accounting for the links between HRF and this behavior. To date, so far as we are aware, no research has examined the relationships between the six motivations underlined by SDT and the promotion and prevention foci. However, Lalot, Quiamzade and Zerhouni [14] have examined the interaction effects of intrinsic and extrinsic motives and message framing in terms of promotion versus prevention on the eating behaviors of students (i.e., personal intention to act, willingness to participate in an online program, interest for nutrition-related information). The authors showed that prevention focus framing worked best to promote nutrition behaviors for participants who reported higher extrinsic motives. However, regarding intrinsic motives, it was more difficult for the authors to draw a clear prediction. These motives, which rely by definition on an internal drive towards action, lead one to expect a reduced or even null impact of the context (promotion or prevention). Furthermore, two studies [15, 16] have investigated the links between regulatory foci and basic psychological needs for autonomy (feeling of choice and of being the initiator of one’s actions), competence (feeling that one interacts efficiently with one’s environment) and relatedness (feeling accepted and recognized by significant others) which are the theoretical foundations of SDT [17]. In particular, Hui et al. [15] examined how chronic or induced promotion and prevention foci can affect the salience of basic needs when individuals evaluate relationship well-being. These authors concluded that promotion-focused individuals judged satisfaction of the need for autonomy as more relevant to evaluate relationship well-being than prevention-focused individuals. In the same vein, Vaughn [16] examined how induced promotion and prevention foci can affect subjective support for the basic needs and how, reciprocally, support for these needs can affect subjective labeling of experiences as promotion- or prevention-focused. People recalled more support for autonomy, competence and relatedness needs in promotion conditions compared with prevention conditions, and experiences of higher need support are more likely to be labeled as promotion-focused rather than prevention-focused.

In line with these previous works, the overall purpose of this paper is to associate for the first time in the literature the RFT and SDT to better understand the dynamics of PA practice in a health context. First, this research proposes to explore the patterns of association of health promotion and prevention foci with the six forms of motivation underlined by SDT (Study 1). Secondly, the indirect associations between HRF and PA behavior through each motivation will be examined (Study 2).

Promotion focus being favorable to support for the needs of autonomy, competence and relatedness, all of which are conceived as the “essential nutriments” of the development of self-determination [17], we hypothesized that this focus in a health context would be positively related with more self-determined forms of motivation for PA (i.e., intrinsic motivation, integrated regulation, and identified regulation). Conversely, prevention focus being less favorable to the support of these needs, this focus in a health context should be related with the less self-determined forms of motivation for PA (i.e., introjected regulation and external regulation) and amotivation. Considering the positive associations between self-determined forms of motivation and PA [13], we further hypothesized that health promotion focus would be indirectly associated with PA through intrinsic motivation, integrated regulation, and identified regulation. On the other hand, considering the predominantly negative associations reported in the literature between controlled forms of motivation and PA [13], we hypothesized that health prevention focus could be negatively related with PA through introjected regulation, external regulation, and amotivation.

## STUDY 1

### Objective

This first cross-sectional study aims to explore for the first time in the literature the patterns of association of health promotion and prevention foci with the six forms of motivation for physical activity (intrinsic motivation, integrated regulation, identified regulation, introjected regulation, external regulation, and amotivation).

### Method

#### Participants

To be eligible for the study participants had to be over 18 and able to read French. The sample comprised 603 French volunteer participants (59.2% men) aged from 18 to 69 years (mean age = 43.8, standard deviation = 14.1, median score = 44, skewness value = .03), either healthy (59.7%) or concerned by a health condition (chronic disease or severe illness during the last 12 months). Most of the participants (95.4%) had completed secondary education. Detailed sample characteristics are presented in Table 1.

**Table 1.** Demographic characteristics of the samples – Study 1

|  | N | % |
| --- | --- | --- |
| **Gender** |  |  |
| Men | 357 | 59.2 |
| Women | 246 | 40.8 |
| **Age** |  |  |
| Young (18-35) | 212 | 35.2 |
| Middle Aged (36-54) | 221 | 36.7 |
| Old (55 and more) | 170 | 28.2 |
| **Health status** |  |  |
| Healthy | 360 | 59.7 |
| Unhealthy* | 243 | 40.3 |
| Chronic joint or back problems | 89 | 14.8 |
| Migraine or chronic headache | 63 | 10.4 |
| Chronic heart disease | 63 | 10.4 |
| Arthritis | 45 | 7.5 |
| Diabetes | 39 | 6.5 |
| Serious dermatologic disorders | 27 | 4.5 |
| Kidney disease | 23 | 3.8 |
| Pulmonary emphysema | 17 | 2.8 |
| Thyroid and gland disorders | 13 | 2.2 |
| Cancer | 12 | 2 |
| Leg ulcer | 10 | 1.7 |
| Asthma or chronic bronchitis | 7 | 1.2 |
| Liver disorder or gallstones | 5 | 0.8 |
| Arterial hypertension | 4 | 0.7 |
| Stroke | 3 | 0.5 |
| Stomach ulcer | 3 | 0.5 |
| Epilepsy | 1 | 0.2 |
| Sclerosis | 1 | 0.2 |
| **Education** |  |  |
| Primary school or lower | 9 | 1.5 |
| Middle school | 92 | 15.3 |
| High school | 121 | 20.1 |
| University | 381 | 63.2 |

\* Unhealthy participants were those which report currently suffered from diseases in the last 12 months. These participants could suffer from one or several diseases.

#### Procedure

Data were collected in June 2017 via a cross-sectional online self-report survey. Participants were recruited via a national online research panel (Dynata, https://www.dynata.com, ISO 20252:2019). Participants were invited by email to participate in an online study. By clicking on the hyperlink provided, they were directed to a secure webpage and were then told they would be taking part in a study on health motivations linked to the practice of a PA. The survey content (instructions, questionnaires) was identical for all participants. Participants were instructed to complete the survey individually in a quiet environment, to be well focused and not to be disturbed for ten minutes. As an incentive, they received points that allowed them to win gift cards. All participants were treated in accordance with the ethical requirements of the Declaration of Helsinki and the French Psychological Society with respect to consent, confidentiality, and anonymity of the answers. Prior to data collection, all participants signed an informed consent form. They were informed of the goal of the study and of their right to stop their participation at any time. The responses were anonymous, as the individuals were only identified by the day and time of completion of the questionnaire. Prior data collection, the study was approved by the CNIL (n°1545711).

#### Measures

##### HRF

Gomez et al.’s [8] French scale was used to assess participants’ HRF. The questionnaire is composed of five items assessing health promotion focus and three items assessing health prevention focus presented in a random order. Participants responded on a scale from 1 = “*strongly disagree*” to 7 = “*strongly agree*”. A Confirmatory Factor Analysis (CFA) performed on the covariance matrix with a maximum likelihood estimation showed that the two-factor model provided a slightly weak fit to the data: *χ*^2^(19) = 124.9; *χ*^2^/*df* = 6.6; CFI = .95; TLI = .93; RMSEA = .10 [90%CI .08-.12]; SRMR = .06. One prevention item (i.e., “*When I implement a health behavior, it’s because I want to protect myself from getting sick*”) exhibited high modification indices and moderate loadings on both factors. These results were similar to those obtained by Schmalbach et al. (2017) and Laroche et al. (in press). In line with the procedure of these studies, a model excluding this item was tested, showing excellent fit indices: *χ*^2^(13) = 36.8; *χ*^2^/*df* = 2.8; CFI = .99; TLI = .98; RMSEA = .06 [90%CI .04 - .08]; SRMR = .03. Internal consistency was satisfactory for the promotion (α = .89) and two-item prevention (α = .77) subscales.

##### Motivations for PA

Boiché, Gourlan, Trouilloud and Sarrazin’s [18] French “*Echelle de Motivation envers l’Activité Physique en contexte de Santé*” (EMAPS) was used to assess participants’ motivations for PA. This questionnaire contains six three-item subscales assessing intrinsic motivation, integrated regulation, identified regulation, introjected regulation, external regulation, and amotivation. Participants responded on a scale from 1 = “*strongly disagree*” to 7 = “*strongly agree*”. A CFA performed on the covariance matrix with a maximum likelihood estimation showed that the six-factor model provided excellent fit to the data: *χ*^2^(120) = 412.4; *χ*^2^/*df* = 3.4; CFI = .97; TLI = .96; RMSEA = .06 [90%CI .06 - .07]; SRMR = .04). Values of internal consistency of the subscales were all satisfactory, ranging from .81 to .91.

#### Data analysis

The descriptive statistics and correlations of the variables were first examined. Then, for each measure (i.e., HRF and motivations for PA, respectively), measurement invariance was examined for gender (i.e., men vs. women), age (young vs. middle-aged and middle-aged vs. older), and health status (healthy vs. unhealthy). A model in which all parameters are freely estimated (configural invariance) was compared with a model in which all factor loadings were constrained to be invariant across groups (i.e., weak or metric invariance). After invariance was verified, models evaluating the contribution of HRF on each form of motivation were tested within each group. Analyses were performed using structural equation modeling in the lavaan package of the R software [19].

### Results

#### Descriptive statistics and correlations

Means, standard deviations and Pearson’s correlation coefficients are presented in Table 2.

**Table 2.**
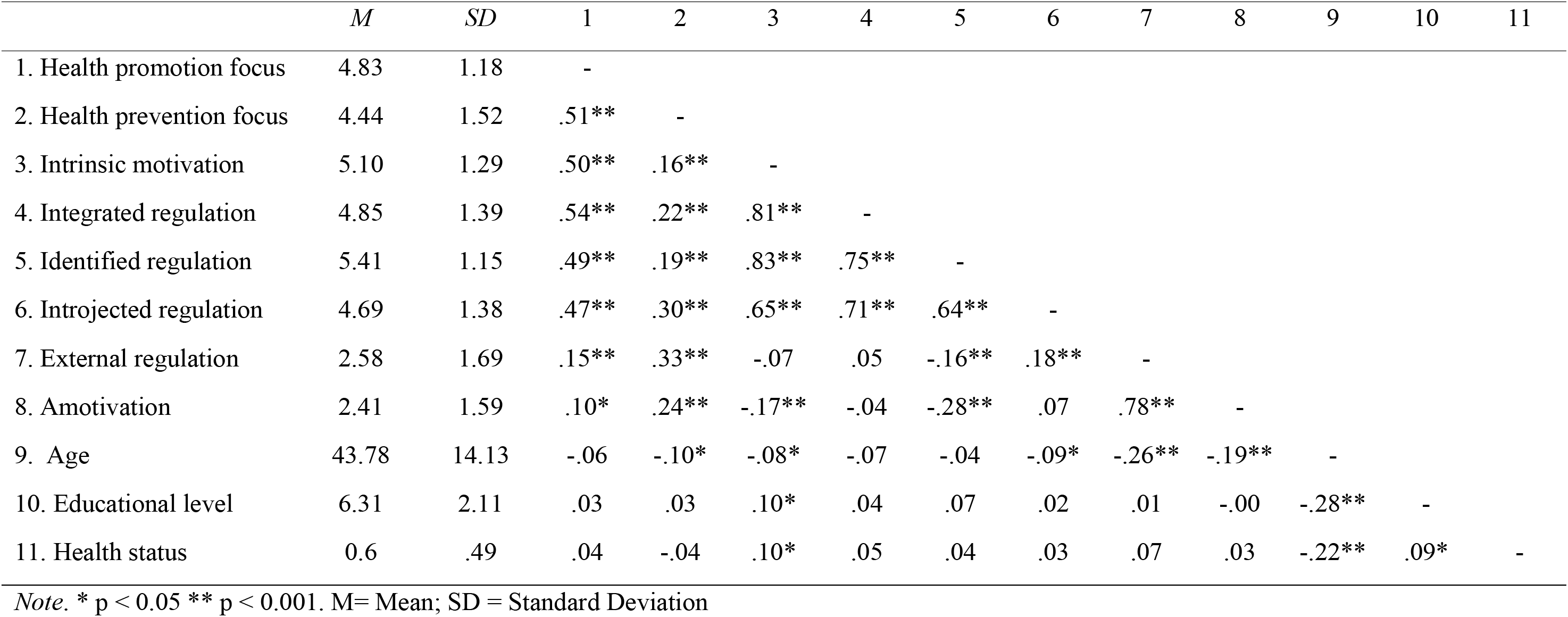
Descriptive statistics and matrix of Pearson r correlation coefficients among the variables – Study 1 (N = 603)

#### Structural equation modeling

All measurements were invariant across gender, age, and health-status groups (i.e., CFI values change < 0.002, [20]; RMSEA value change < 0.015, [21]). Fit indices of the models and beta coefficients in the whole sample and in each group of participants are shown in Table 3. All models provided a good fit to the data. The main results showed that in the whole sample and regardless of gender, age, and health-status groups, health promotion focus was positively related with intrinsic motivation (.84 < β < .62, p <.001), and with integrated regulation (.62 < β < .77, p <.001), identified regulation (.63 < β < .72, p <.001), and introjected regulation (.37 < β < .66, p <.001), whereas health prevention focus was positively related with external regulation (.32 < β < .55, p <.001) and amotivation (.24 < β < .48, p <.001). Furthermore, introjected regulation was not associated with health prevention focus in the whole sample (β = .05, ns.). However, this regulation was positively related with health prevention focus in the group of healthy participants (β = .22, p < .01) and negatively in the group of participants concerned by a health condition (β = −19, p < .01).

**Table 3.** Results of the hypothesized model in the whole sample and in groups of participants – Study 1 (N = 603)

| | $\chi^2$ | *df* | $\chi^2/df$ | CFI | TLI | RMSEA | 90%CI | $\beta_{\text{promotion}}$ | $\beta_{\text{prevention}}$ | R<sup>2</sup> |
| --- | --- | --- | --- | --- | --- | --- | --- | --- | --- | --- |
| **Intrinsic Motivation** | **86.3** | **32** | **2.7** | **.98** | **.98** | **.05** | **[.04-.07]** | **.74** | **-.27** | **.37** |
| Women | 80.9 | 32 | 2.5 | .96 | .94 | .08 | [.06-.10] | .68 | -.22 | .35 |
| Men | 63.1 | 32 | 2.0 | .98 | .98 | .05 | [.03-.07] | .78 | -.31 | .38 |
| Young | 56.5 | 32 | 1.8 | .98 | .97 | .06 | [.03-.09] | .84 | -.36 | .38 |
| Middle Aged | 83.6 | 32 | 2.6 | .96 | .94 | .08 | [.06-.11] | .73 | -.27 | .37 |
| Older | 44.1 | 32 | 1.4 | .99 | .98 | .05 | [.00-.08] | .62 | -.20 | .30 |
| Unhealthy | 67.6 | 32 | 2.1 | .97 | .96 | .07 | [.05-.10] | .72 | -.33 | .34 |
| Healthy | 88.6 | 32 | 2.8 | .97 | .96 | .07 | [.05-.09] | .72 | -.20 | .38 |
| **Integrated Regulation** | **70.6** | **32** | **2.2** | **.99** | **.98** | **.05** | **[.03-.06]** | **.72** | **-.18** | **.39** |
| Women | 69.8 | 32 | 2.2 | .97 | .96 | .07 | [.05-.09] | .66 | -.15 *ns.* | .35 |
| Men | 37.3 | 32 | 1.2 | .99 | .99 | .02 | [.00-.05] | .76 | -.21 | .41 |
| Young | 48.7 | 32 | 1.5 | .99 | .98 | .05 | [.02-.08] | .74 | -.20 *ns.* | .37 |
| Middle Aged | 56.7 | 32 | 1.8 | .98 | .97 | .06 | [.03-.08] | .77 | -.23 | .44 |
| Older | 55.1 | 32 | 1.7 | .99 | .97 | .07 | [.03-.09] | .62 | -.12 *ns.* | .33 |
| Unhealthy | 75.4 | 32 | 2.4 | .97 | .95 | .08 | [.05-.10] | .66 | -.21 | .31 |
| Healthy | 62.0 | 32 | 1.9 | .99 | .98 | .05 | [.03-.07] | .74 | -.13 *ns.* | .44 |
| **Identified Regulation** | **89.1** | **32** | **2.8** | **.98** | **.98** | **.05** | **[.04-.07]** | **.69** | **-.22** | **.33** |
| Women | 74.2 | 32 | 2.3 | .97 | .95 | .07 | [.05-.10] | .64 | -.21 | .31 |
| Men | 64.6 | 32 | 2.0 | .98 | .98 | .05 | [.03-.07] | .72 | -.23 | .34 |
| Young | 64.7 | 32 | 2.0 | .97 | .96 | .07 | [.05-.09] | .72 | -.66 | .30 |
| Middle Aged | 65.4 | 32 | 2.0 | .97 | .96 | .07 | [.05-.09] | .72 | -.25 | .36 |
| Older | 44.2 | 32 | 1.4 | .99 | .98 | .05 | [.00-.08] | .63 | -.13 *ns.* | .33 |
| Unhealthy | 70.9 | 32 | 2.2 | .97 | .95 | .07 | [.05-.09] | .63 | -.25 | .28 |
| Healthy | 74.4 | 32 | 2.3 | .97 | .97 | .06 | [.04-.08] | .71 | -.18 | .37 |

**Table 3.** (Continued)
| | $\chi^2$ | $df$ | $\chi^2/df$ | CFI | TLI | RMSEA | 90%CI | $\beta_{\text{promotion}}$ | $\beta_{\text{prevention}}$ | R<sup>2</sup> |
| --- | --- | --- | --- | --- | --- | --- | --- | --- | --- | --- |
| **Introjected Regulation** | **88.1** | **32** | **2.8** | **.98** | **.97** | **.05** | **[.04-.07]** | **.53** | **.05 *ns.*** | **.31** |
| Women | 82.0 | 32 | 2.6 | .96 | .94 | .08 | [.05-.10] | .37 | .16 *ns.* | .22 |
| Men | 53.1 | 32 | 1.7 | .99 | .98 | .04 | [.02-.06] | .66 | -.07 *ns.* | .37 |
| Young | 56.2 | 32 | 1.8 | .99 | .97 | .06 | [.03-.09] | .41 | .16 *ns.* | .29 |
| Middle Aged | 69.5 | 32 | 2.2 | .97 | .95 | .07 | [.05-.10] | .63 | -.05 *ns.* | .37 |
| Older | 54.8 | 32 | 1.7 | .97 | .96 | .07 | [.03-.09] | .49 | .05 *ns.* | .27 |
| Unhealthy | 84.7 | 32 | 2.6 | .95 | .93 | .08 | [.06-.10] | .62 | -.19 | .29 |
| Healthy | 63.5 | 32 | 2.0 | .98 | .98 | .05 | [.03-.07] | .43 | .22 | .35 |
| **External Regulation** | **50.3** | **32** | **1.6** | **.98** | **.98** | **.06** | **[.04-.07]** | **-.13** | **.45** | **.15** |
| Women | 82.0 | 32 | 2.6 | .96 | .95 | .08 | [.06-.10] | -.18 | .45 | .14 |
| Men | 55.6 | 32 | 1.7 | .99 | .98 | .05 | [.02-.07] | -.09 *ns.* | .45 | .16 |
| Young | 73.9 | 32 | 2.3 | .97 | .96 | .08 | [.06-.10] | -.10 *ns.* | .51 | .19 |
| Middle Aged | 58.3 | 32 | 1.8 | .98 | .97 | .06 | [.04-.09] | -.21 | .46 | .14 |
| Older | 54.7 | 32 | 1.7 | .97 | .96 | .07 | [.03-.09] | -.12 *ns.* | .32 | .08 |
| Unhealthy | 80.5 | 32 | 2.5 | .96 | .95 | .08 | [.06-.10] | -.16 *ns.* | .34 | .08 |
| Healthy | 82.6 | 32 | 2.6 | .98 | .97 | .07 | [.05-.08] | -.13 *ns.* | .55 | .23 |
| **Amotivation** | **68.6** | **32** | **2.1** | **.99** | **.89** | **.04** | **[.03-.08]** | **-.13** | **.38** | **.10** |
| Women | 64.6 | 32 | 2.0 | .98 | .97 | .06 | [.04-.09] | -.13 *ns.* | .34 | .09 |
| Men | 51.7 | 32 | 1.6 | .99 | .99 | .04 | [.02-.06] | -.13 *ns.* | .42 | .12 |
| Young | 53.9 | 32 | 1.7 | .98 | .98 | .06 | [.03-.08] | .09 *ns.* | .42 | .21 |
| Middle Aged | 57.3 | 32 | 1.8 | .98 | .97 | .06 | [.03-.08] | -.31 | .34 | .09 |
| Older | 39.8 | 32 | 1.2 | .99 | .99 | .04 | [.00-.07] | .17 *ns.* | .27 | .06 |
| Unhealthy | 78.9 | 32 | 2.5 | .96 | .95 | .08 | [.06-.10] | -.08 *ns.* | .24 | .04 |
| Healthy | 55.0 | 32 | 1.7 | .99 | .98 | .05 | [.02-.06] | -.19 | .49 | .16 |

## STUDY 2

### Objective

After examining in a first exploratory study the patterns of association of health promotion and prevention foci with the six forms of motivation underlined by SDT, we aimed in this second cross-sectional study to investigate to what extent these six forms of motivation could play a mediating role in the links between HRF and PA behavior.

### Method

#### Participants

As in Study 1, to be eligible for the study participants had to be over 18 and able to read French. A total of 395 French volunteer participants (59.7% men) aged from 19 to 71 years (mean age = 41.4; standard deviation = 14.7; median score = 40, skewness value = .20), either healthy (57%) or concerned by a health condition (chronic disease or severe illness during the last 12 months) took part in the study. Most of the participants (98.6%) had completed secondary education. Detailed sample characteristics are presented in Table 4.

**Table 4.** Demographic characteristics of the samples – Study 2

|  | N | % |
| --- | --- | --- |
| **Gender** |  |  |
| Men | 236 | 59.7 |
| Women | 159 | 40.3 |
| **Age** |  |  |
| Young (18-35) | 157 | 39.7 |
| Middle Aged (36-54) | 146 | 36.9 |
| Old (55 and more) | 92 | 23.3 |
| **Health status** |  |  |
| Healthy | 225 | 57 |
| Unhealthy* | 170 | 43 |
| Chronic joint or back problems | 38 | 9.5 |
| Migraine or chronic headache | 34 | 8.5 |
| Chronic heart disease | 9 | 2.6 |
| Arthritis | 31 | 7.8 |
| Diabetes | 1 | 0.3 |
| Serious dermatologic disorders | 43 | 10.8 |
| Kidney disease | 4 | 1 |
| Pulmonary emphysema | 12 | 3 |
| Thyroid and gland disorders | 0 | 0 |
| Cancer | 12 | 3 |
| Leg ulcer | 13 | 3.3 |
| Asthma or chronic bronchitis | 23 | 3.8 |
| Liver disorder or gallstones | 6 | 0.8 |
| Arterial hypertension | 39 | 9.8 |
| Stroke | 19 | 4.8 |
| Stomach ulcer | 4 | 1.5 |
| Epilepsy | 1 | 0.3 |
| Sclerosis | 4 | 1 |
| **Education** |  |  |
| Primary school or lower | 18 | 4.6 |
| Middle school | 60 | 15.2 |
| High school | 91 | 23 |
| University | 226 | 57.2 |

\* Unhealthy participants were those which report currently suffered from diseases in the last 12 months. These participants could suffer from one or several diseases.

#### Procedure

Data were collected in April 2018 via a cross-sectional online self-report survey. As in Study 1, participants were recruited via the national online research panel Dynata (https://www.dynata.com, ISO 20252:2019). The procedure and instructions were similar to those of Study 1. Prior data collection, the study was also approved by the CNIL (n°1545711).

#### Measures

##### HRF

A modified version of Gomez et al.’s [8] scale was used to assess participants’ HRF. The prevention item excluded in Study 1 was slightly rephrased (“ *When I think about my health, I often imagine diseases that I could have*”). A CFA performed on the covariance matrix with a maximum likelihood estimation showed that the two-factor model provided a good fit to the data: *χ*^2^(19) = 55; *χ*^2^/*df* = 2.9; CFI = .98; TLI = .97; RMSEA = .07 [90%CI .05 - .09]; SRMR = .05. Internal consistency was satisfactory for both promotion (α = .87) and prevention (α = .87) subscales.

##### Motivations for PA

As in Study 1, Boiché et al.’s [18] EMAPS was used to assess participants’ motivations for PA. A CFA performed on the covariance matrix with a maximum likelihood estimation showed that the six-factor model provided a good fit to the data: *χ*^2^(120) = 362.8; *χ*^2^/*df* = 3.02; CFI = .96; TLI = .95; RMSEA = .07 [90%CI .06 - .08]; SRMR = .05. Internal consistency was satisfactory for each subscale, ranging from .79 to .92.

##### Self-reported PA

The French long form (27 items) of the International Physical Activity Questionnaire [22] was used to assess self-reported PA. This questionnaire is appropriate for both healthy adults [23] and patients with chronic diseases [24]. Participants were asked to rate the frequency and duration of vigorous, moderate, and walking activity across four domains of living (work, transport, chores, and leisure) performed during the last seven days. Following standard procedures (available for download at http://www.ipaq.ki.se), a total weekly PA was calculated. Because the distribution of total PA scores was strongly skewed, in line with previous studies [25] a logarithmic transformation (log) was used to improve the normality of the distribution.

#### Data analysis

The descriptive statistics and correlations between variables were first examined. Then, six models were successively run to test indirect associations of HRF with self-reported PA through each form of motivation for PA. All models included age, gender, and health status as control variables. Analyses were performed using structural equation modeling in the lavaan package of the R software. Finally, using the process package [26] in the SPSS software, a bootstrapping method resample set at 5000 samples with bias-corrected 95% Confidence Intervals (CI) was employed to test the significance of indirect effects.

### Results

#### Descriptive statistics and correlations

Means, standard deviations and Pearson’s correlation coefficients are presented in Table 5.

**Table 5.** Descriptive statistics and matrix of Pearson r correlation coefficients among the variables – Study 2 (N = 395).

|  | *M* | *SD* | 1 | 2 | 3 | 4 | 5 | 6 | 7 | 8 | 9 | 10 | 11 | 12 |
| --- | --- | --- | --- | --- | --- | --- | --- | --- | --- | --- | --- | --- | --- | --- |
| 1. Health promotion focus | 4.71 | 1.24 | - |  |  |  |  |  |  |  |  |  |  |  |
| 2. Health prevention focus | 4.23 | 1.56 | .45** | - |  |  |  |  |  |  |  |  |  |  |
| 3. Intrinsic motivation | 4.67 | 1.48 | .44** | .16** | - |  |  |  |  |  |  |  |  |  |
| 4. Integrated regulation | 4.48 | 1.61 | .47** | .21** | .86** | - |  |  |  |  |  |  |  |  |
| 5. Identified regulation | 5.02 | 1.41 | .40** | .12* | .84** | .82** | - |  |  |  |  |  |  |  |
| 6. Introjected regulation | 4.43 | 1.47 | .41** | .26** | .79** | .84** | .76** | - |  |  |  |  |  |  |
| 7. External regulation | 2.84 | 1.78 | .20** | .33** | .12* | .19** | .02 | .29** | - |  |  |  |  |  |
| 8. Amotivation | 2.88 | 1.65 | .16** | .33** | -.13* | -.05 | -.21** | .02 | .69** | - |  |  |  |  |
| 9. Age | 41.38 | 14.65 | -.05 | -.17** | -.12* | -.09 | .00 | -.16** | -.25** | -.17** | - |  |  |  |
| 10. Educational level | 5.88 | 2.07 | .03 | .04 | .17** | .16** | .16** | .15** | .11* | .09 | -.28** | - |  |  |
| 11. Health status | .57 | .50 | -.07 | -.07 | .09 | .06 | .05 | .06 | -.07 | -.09 | -.13* | .05 | - |  |
| 12. Self-reported PA (log) | 3.83 | .80 | .16** | .08 | .24** | .28** | .21** | .24** | .16* | .01 | .03 | -.14** | -.06 | - |

*Note.* \* $p < 0.05$ \*\* $p < 0.001$ . M= Mean; SD = Standard Deviation

#### Structural equation modeling

Fit indices, beta coefficients and bootstrapped CI for each model are shown in Table 6. All models provided a good fit to the data. While controlling for age, gender, and health status, health promotion focus was positively related with intrinsic motivation (β =.44, p < .001), integrated regulation (β =.47, p < .001), identified regulation (β =.40, p < .001), and introjected regulation (β =.41, p < .001), which were all positively related with PA (.21 < β < .28, p < .01). Bootstrapping analyses indicated that these four indirect associations were significant (95% bootstrapped CI [.02 to .11] for intrinsic motivation, 95% bootstrapped CI [.00 to .08] for integrated regulation, 95% bootstrapped CI [.00 to .09] for identified regulation, and 95% bootstrapped CI [.04 to .12] for introjected regulation). On the other hand, health prevention focus was positively related both with external regulation (β = .31, p < .001), which was positively related with PA (β = .15, p < .05), and with amotivation (β = .32, p < .001), which was not related with PA (β = -.03, *ns*.). Bootstrapping analyses indicated a significant indirect association of health prevention focus with PA through external regulation (95% bootstrapped CI [.00 to .12]). Significant indirect relations are illustrated in Fig 1.

**Fig 1.**
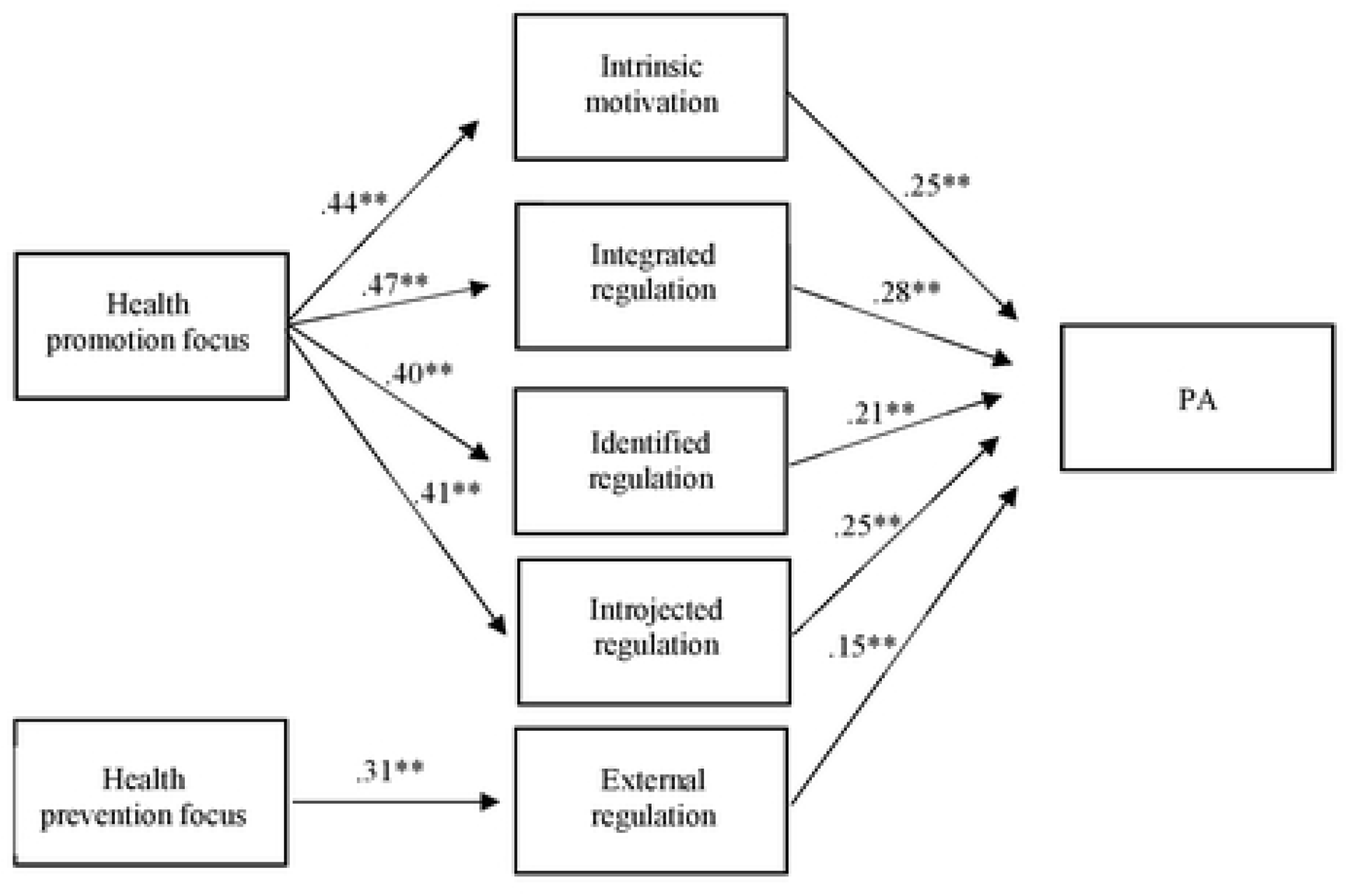
Summary of significant indirect relations between HRF with PA through motivations while controlling for gender, age and health status. *Note*. Path values arc standardized β coefficients. \*\**p* ≤ 0.01.

**Table 6.** Summary of the fit, pathways and bootstrapped CI of each of the six-process models of PA – Study 2 (N = 395)

| | $\beta$ | bootstrapped 95% CI | $R^2$ | $\chi^2$ | $df$ | $\chi^2/df$ | CFI | TLI | RMSEA | 90%CI |
| --- | --- | --- | --- | --- | --- | --- | --- | --- | --- | --- |
| **Intrinsic motivation model** |  |  | **.08** | **13.4** | **5** | **2.7** | **.93** | **.84** | **.07** | **[.02-.11]** |
| Health promotion focus → Intrinsic motivation | .44 |  |  |  |  |  |  |  |  |  |
| Intrinsic motivation → PA | .25 |  |  |  |  |  |  |  |  |  |
| Health promotion focus → Intrinsic motivation → PA |  | [.02; .11] |  |  |  |  |  |  |  |  |
| **Integrated regulation model** |  |  | **.09** | **6** | **5** | **1.2** | **.99** | **.98** | **.02** | **[.00-.08]** |
| Health promotion focus → Integrated regulation | .47 |  |  |  |  |  |  |  |  |  |
| Integrated regulation → PA | .28 |  |  |  |  |  |  |  |  |  |
| Health promotion focus → Integrated regulation → PA |  | [.00; .08] |  |  |  |  |  |  |  |  |
| **Identified regulation model** |  |  | **.06** | **8.3** | **5** | **1.7** | **.96** | **.92** | **.04** | **[.00-.09]** |
| Health promotion focus → Identified motivation | .40 |  |  |  |  |  |  |  |  |  |
| Identified regulation → PA | .21 |  |  |  |  |  |  |  |  |  |
| Health promotion focus → Identified regulation → PA |  | [.00; .09] |  |  |  |  |  |  |  |  |
| **Introjected regulation model** |  |  | **.07** | **17.5** | **5** | **3.5** | **.88** | **.74** | **.08** | **[.04-.12]** |
| Health promotion focus → Introjected regulation | .41 |  |  |  |  |  |  |  |  |  |
| Introjected regulation → PA | .25 |  |  |  |  |  |  |  |  |  |
| Health promotion focus → Introjected regulation → PA |  | [.04; .12] |  |  |  |  |  |  |  |  |
| **External regulation model** |  |  | **.05** | **7.3** | **3** | **2.4** | **.95** | **.83** | **.06** | **[.00-.12]** |
| Health prevention focus → External regulation | .31 |  |  |  |  |  |  |  |  |  |
| External regulation → PA | .15 |  |  |  |  |  |  |  |  |  |
| Health prevention focus → External regulation → PA |  | [.00; .12] |  |  |  |  |  |  |  |  |
| **Amotivation model** |  |  | **.03** | **6** | **2** | **3** | **.94** | **.68** | **.07** | **[.00-.14]** |
| Health prevention focus → Amotivation | .32 |  |  |  |  |  |  |  |  |  |
| Amotivation → PA | -0.3 *ns.* |  |  |  |  |  |  |  |  |  |
| Health prevention focus → Amotivation → PA |  | [-.03; .02] *ns.* |  |  |  |  |  |  |  |  |

Note: in each model gender, age and health status effects were controlled

## DISCUSSION

The present research investigated for the first time in the literature the empirical links between HRF with each form of motivation for PA underlined by SDT, and the indirect associations between HRF and PA practice through each of these motivations for PA. Our results strongly support our hypothesis that concerns about attaining health-related gains are favorable to autonomy and the development of self-determined motivations toward PA (intrinsic motivation, integrated regulation, and identified regulation), whereas concerns about avoiding health-related losses are associated with a practice based on a feeling of external pressures (external regulation) and an incapacity to value the activity or its outcomes (amotivation). These links showed robust support across the two studies and analyses conducted in sub-groups of participants indicated that they were consistent across gender, age, and health status.

However, our results showed that introjected regulation was not associated with HRF in the expected direction. Indeed, in both studies, introjected regulation was positively related with health promotion focus, but unrelated with health prevention focus. These unexpected results could be attributed to the fact that introjected regulation in a health-related PA context is strongly associated with self-determined forms of motivation [18]. Moreover, while the positive link between health promotion focus and introjected regulation was invariant in Study 1 whatever the characteristics of the participants, the link between health prevention focus and this motivation was completely opposite depending on the health status of participants. Among unhealthy adults, health prevention focus could thus constitute an obstacle to the first step of the internalization process [17] and generate a practice exclusively driven by external pressures. This result illustrates the importance of considering illness experience to understand the motivational issues surrounding PA practice.

Regarding the indirect associations between HRF and PA through motivations, the results of Study 2 strongly support the hypothesis that health promotion focus is positively associated with PA through more self-determined motivation (intrinsic motivation, integrated regulation, and identified regulation). However, they partially support the hypothesis that health prevention focus is positively associated with PA through more controlled motivation (introjected regulation, external regulation) and amotivation. Indeed, only external regulation was found to be a relevant candidate to explain the link between health prevention focus and PA. In addition, unexpectedly, health prevention focus was found to be *positively* related with PA through this regulation.

On the one hand, the indirect positive relationship between health prevention focus and PA suggests that health prevention focus is not systematically detrimental to the practice of a PA. This finding is not in line with Laroche et al. [10], which evidences that among sport practitioners, health prevention focus is negatively related with amount of sports practice. This contrasting result might be explained by the fact that our study is interested in general PA and not specifically with sport practice. The idea of practicing at least a minimum of PA is now generally widespread in medical discourse and in public health recommendations. PA might therefore be perceived as a more “medicalized” practice than sport and thus be more compatible with individual preoccupations centered on avoiding illness. Moreover, the fact that we worked on a general population and not exclusively with sport practitioners could also help to explain this result. Indeed, Pfeffer [27], based on the Regulatory Fit Theory [28], showed that among non-sport practitioners, prevention-oriented participants have a greater intention to practice when they are confronted with a health communication highlighting PA as a means of avoiding health problems. These results thus evidence that among non-sport practitioners, PA is not incompatible with a focus on avoiding health problems. This question would nonetheless merit further exploration in subsequent studies specifically comparing the link between health prevention focus and PA in these two populations (sport practitioners *vs*. non-sport practitioners).

On the other hand, the positive link between external regulation and PA suggests that external regulation is not systematically detrimental for PA. This result does not support the majority of works in past literature [13]. However, it is in line with two other studies reporting that among people with a high level of practice, some display a motivational profile characterized by high levels of external regulation [29, 30]. It thus appears necessary to continue exploring to what extent individuals with a high level of external regulation are physically active, depending on their health prevention focus. Moreover, considering that some works suggest that external regulation has a beneficial effect on PA practice only in the short term [13], it would be interesting to examine, in a complementary approach to this cross-sectional study, the nature of the indirect link between health prevention focus and PA through external regulation over time.

Despite these unexpected results, all the data of these two exploratory studies nonetheless largely support our hypotheses. In this sense, these results on the link between the RFT and SDT frameworks, which has been hitherto very little studied, may encourage researchers to pursue analysis of this theoretical association so as to study PA practice in a health context. Moreover, our results contribute to the literature at three levels. First, they confirm the interest of associating these two theoretical models in order to study a behavior. In this respect, they complement both the previous work of Lalot et al. [14], who combined these two models to study nutrition habits, and the works of Vaughn [16] and Hui et al. [15], who associated these two theoretical frameworks with another concept underlined by SDT (i.e., basic needs), in contexts other than health. Secondly, they improve understanding of the process through which health promotion focus is related with PA and thus complement the work of Laroche et al. [10] showing a mediator in this relationship (i.e., SOC strategy). Thirdly, for the first time in the literature they provide a better understanding of the process through which health prevention focus is related with PA.

These results also point to practical steps that can be taken to better promote PA. They suggest that health communication and the coaching arguments in health apps focusing on health improvement (e.g., “*PA is good for your health*”) should encourage PA practice by favoring the pursuit of enjoyment in the activity and the acknowledgment of its usefulness. On the other hand, health messages and coaching arguments in health apps focusing on avoidance of health-related problems (e.g., “*PA protects against health threats*”) should favor practice motivated by external pressures (e.g., fear of disease, fear of reproaches from certain people such as doctors or family) which are also favorable to practice. However, the works based on SDT suggest that practice motivated by external pressures is not beneficial for individuals’ fulfillment and well-being [11] and only favor PA practice in the short term [13]. Therefore, this finding nonetheless casts doubt on the long-term efficacy of health messages focused on avoidance of health-related problems.

